# A control-validated pan-proteome deep-learning pipeline nominates GPR35 as a candidate target of the orphan bacterial metabolite ligiamycin A

**DOI:** 10.64898/2026.07.01.735807

**Authors:** James K. Martin

**Affiliations:** Department of Medical Education and Scholarship, Rowan-Virtua School of Osteopathic Medicine, Stratford, NJ, USA

## Abstract

Most microbial natural products with documented bioactivity lack an identified molecular target, which limits their development. We present an open, control-validated computational pipeline for natural-product target hypothesis generation. It combines a pan-proteome deep-learning drug-target interaction (DTI) model (a graph neural-network ligand encoder, an ESM-2 protein language-model encoder, and bidirectional cross-attention) with bias-corrected ranking and control-anchored molecular docking. Applying it to ligiamycin A, a 2022-described Streptomyces/Achromobacter co-culture decalin-amino-maleimide with no reported target, we find that the predicted interactions of the compound are dominated by class-A G-protein-coupled receptors. Using a drug with a known target (losartan) we identify and correct a frequent-hitter bias in the raw model; after correction the standout candidates are uniformly class-A GPCRs, led by the orphan receptor GPR35. Structure-based docking with matched positive and negative controls across three candidates corroborates GPR35 specifically: ligiamycin A scores comparably to the known GPR35 agonist zaprinast at the agonist pocket (-8.1 vs -8.3 kcal/mol; non-binder floor -5.5), whereas FFAR1 is excluded and histamine H2 is inconclusive. We propose GPR35 as a prioritized, experimentally testable target and release the workflow as a reusable tool. The result is a computational hypothesis that requires experimental validation.

## Introduction

Natural products remain a primary source of antibiotics and anticancer leads, yet for the majority the protein target underlying their phenotype is unknown. Experimental target deconvolution (affinity pulldown, thermal proteome profiling, genetic screens) is powerful but slow and resource-intensive. Computational DTI models trained on large bioactivity databases offer rapid triage, but two problems limit their use on genuinely novel chemotypes. First, models trained on imbalanced data over-predict a small set of heavily-measured “frequent-hitter” proteins. Second, single scores are hard to trust without orthogonal evidence. Our pipeline adapts the recently described HitScreen framework for sequence-based virtual screening [1], which targets class-imbalance and annotation/ligand biases in DTI training data, and pairs it with explicit frequent-hitter correction and control-anchored docking. We demonstrate the workflow on ligiamycin A [2], a 2022-described bacterial decalin-amino-maleimide with antibacterial activity and no published binding partner.

**Figure 1.**
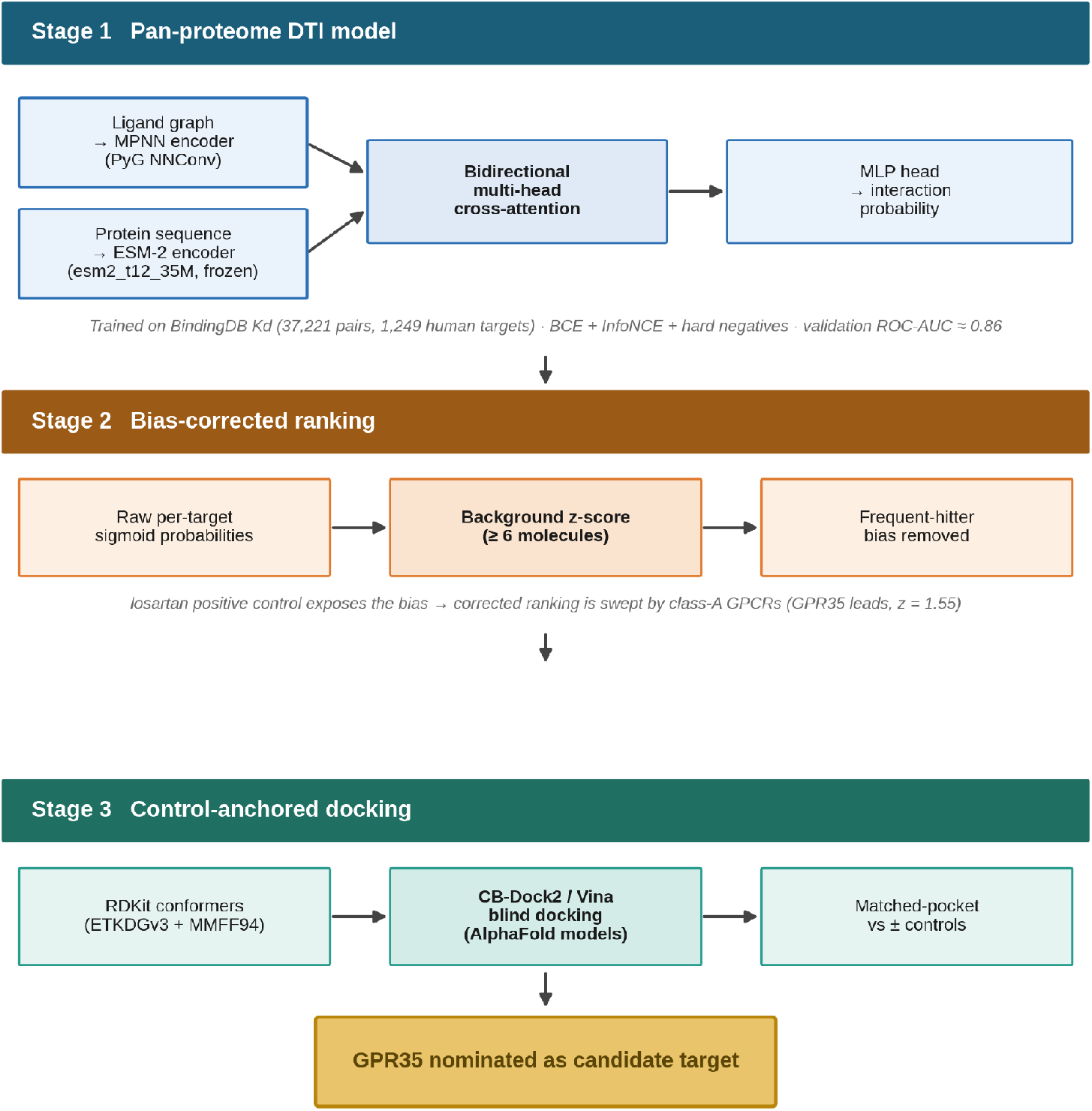
Pipeline schematic. The three stages of the workflow: a pan-proteome DTI model (MPNN ligand encoder, frozen ESM-2 protein encoder, bidirectional cross-attention fusion, MLP head) trained on the BindingDB Kd subset; background z-score correction that removes frequent-hitter bias from the raw ranking; and control-anchored blind docking against AlphaFold models with matched positive and negative controls. The workflow nominates GPR35 as the prioritized candidate target.

## Results

### A pan-proteome DTI model and orphan-compound screen

We trained a DTI model (Methods) on the BindingDB Kd subset [3] (37,221 ligand–target pairs; 1,249 unique human targets) to a validation ROC-AUC of about 0.86. Screening ligiamycin A against all unique targets returned a top-ranked set dominated by class-A GPCRs (neurotensin receptor 2, angiotensin II type-1 receptor, and histamine H4), with a single receptor tyrosine kinase, FLT3, intruding at rank 7 (Figure 2a; Supplementary S1). Of the 1,249 targets, 128 cleared the active threshold (probability ≥ 0.50).

**Figure 2.**
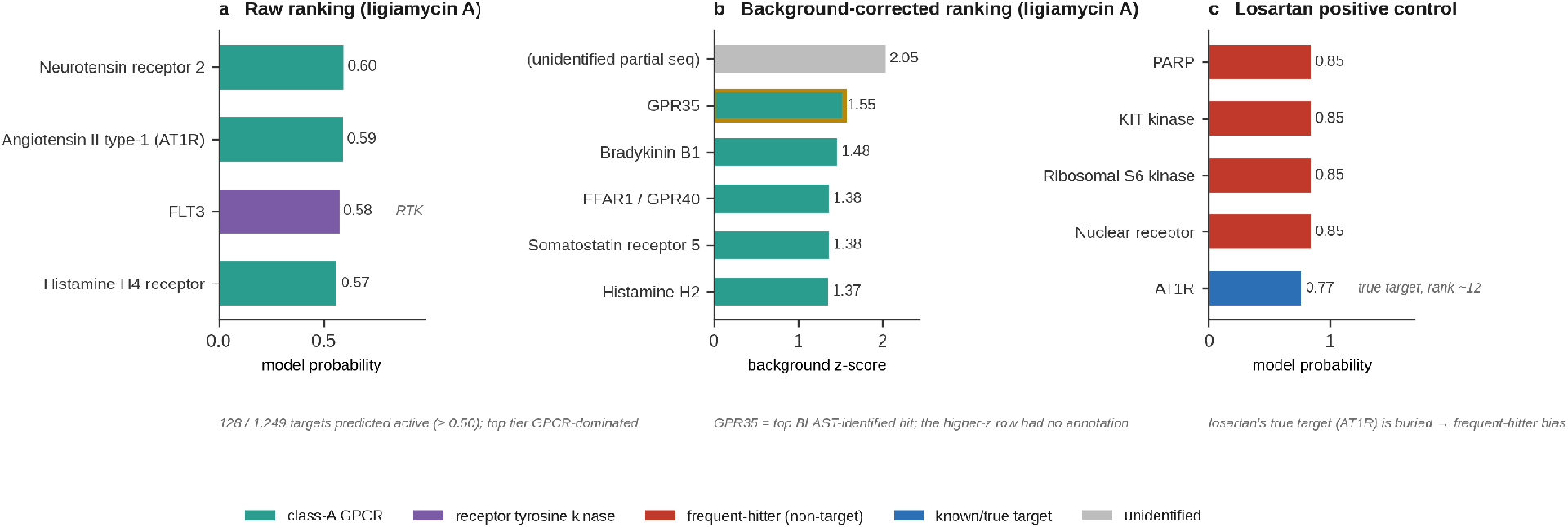
Bias correction reshapes the ranking. (a) Raw model probabilities for the top ligiamycin A targets: the tier is GPCR-dominated, with one receptor tyrosine kinase (FLT3) intruding, and 128 of 1,249 targets clear the active threshold (≥ 0.50). (b) Background-corrected ranking by per-target z-score: the credible top tier is uniformly class-A GPCRs, led by GPR35 at z = 1.55; the single higher-z entry was an unannotatable partial sequence. (c) Losartan positive control: the drug’s true target, AT1R, is buried near rank 12 while heavily-measured frequent-hitter proteins top the list, which is the bias the z-score correction removes. Full tables in Supplementary S1–S3.

### Identifying and correcting frequent-hitter bias

A positive-control screen with losartan, a selective angiotensin AT1R antagonist, failed to rank AT1R first. Instead a fixed set of heavily-measured proteins (a PARP, KIT, ribosomal-S6 kinases, a nuclear receptor) topped the list, as they did for unrelated ligands (Figure 2c; Supplementary S3). We therefore re-ranked targets by a per-target z-score relative to a background of other molecules. After correction, ligiamycin A’s standout targets were uniformly class-A GPCRs: GPR35 (the top BLAST-identified hit, z = 1.55; the single higher-z row was an unannotatable partial sequence), bradykinin B1, FFAR1/GPR40, somatostatin receptor 5, and histamine H2 (Figure 2b; Supplementary S2). Several of these are lipid- or peptide-sensing receptors, consistent with the compound’s lipophilic decalin scaffold.

### Control-anchored docking corroborates GPR35 specifically

For three candidate receptors we performed cavity-detection blind docking [4] (AutoDock Vina [5]) against AlphaFold models [6,7], each with a known-ligand positive control and a glucose negative control, comparing scores at the pocket identified by the positive control (Figure 3; Supplementary S4):

**Figure 3.**
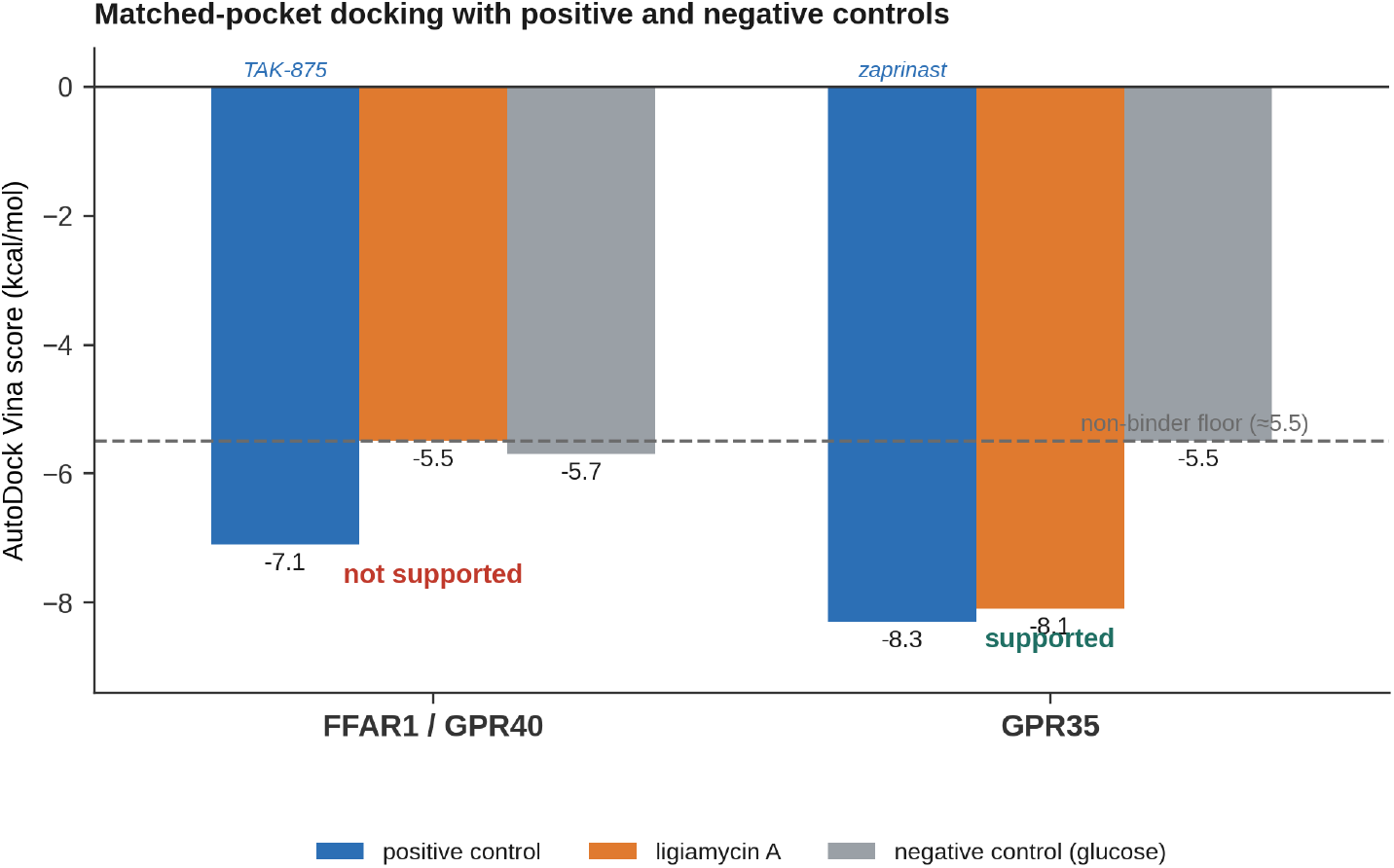
Control-anchored matched-pocket docking. AutoDock Vina scores at the pocket defined by each receptor’s positive control (more negative is stronger binding). The dashed line marks the non-binder floor (≈ −5.5 kcal/mol) set by the glucose negative control. At GPR35, ligiamycin A (−8.1) scores within 0.2 kcal/mol of the known agonist zaprinast (−8.3) and about 2.6 kcal/mol below the floor; at FFAR1 it sits at the floor alongside glucose. The inconclusive histamine H2 test (collapsed apo pocket) is reported in Supplementary S4d. A pose overlay of ligiamycin A and zaprinast in the GPR35 pocket will be added as a final panel from the docking output coordinates.

- **FFAR1:** docking against the FFAR1 crystal structure (4PHU) first gave ligiamycin A a misleadingly favorable −7.8 kcal/mol, but the pose sat on the construct’s T4-lysozyme crystallization fusion rather than the receptor (Supplementary S4a). The fusion-free AlphaFold model removed this artifact: there the native agonist TAK-875 docked correctly (−7.1 kcal/mol at the orthosteric pocket) while ligiamycin A scored like the non-binder glucose (−5.5 vs −5.7). **Not supported**.
- **GPR35:** the agonist zaprinast docked at −8.3 kcal/mol; ligiamycin A scored −8.1 at the same pocket, about 2.6 kcal/mol below the glucose floor (−5.5). Zaprinast and ligiamycin are similar in size, which excludes a size/lipophilicity artifact. **Supported**.

A third receptor, histamine H2, was inconclusive: its positive control (famotidine) scored at the non-binder floor, indicating a collapsed, ligand-free AlphaFold pocket, so ligiamycin A’s favorable score there is uninterpretable. We report H2 in Supplementary S4d and restrict Figure 3 to the two interpretable receptors.

Of three receptors tested, docking supported only GPR35, the same receptor ranked first by the corrected model, and it did so discriminatingly, rejecting FFAR1 and flagging H2 as invalid rather than rubber-stamping every candidate.

## Discussion

Two independent methods (a bias-corrected DTI model and control-anchored docking) converge on GPR35 as the leading candidate target of ligiamycin A. We want to be honest about the scope of the claim. The losartan control shows the model resolves target class (GPCR) more reliably than specific identity, and all evidence here is computational. GPR35 is an understudied, therapeutically interesting receptor, with immuno-metabolic, gastrointestinal, and pain biology and debated endogenous ligands; a defined natural-product chemotype engaging it would be valuable as a chemical probe. The pipeline’s value is as much methodological as specific: explicit frequent-hitter correction and control-anchored docking convert a noisy ranking into a discriminating, falsifiable hypothesis.

### Limitations

This is a computational study. Training was CPU-limited (ROC-AUC ≈ 0.86), and the panel is restricted to BindingDB Kd targets, so compounds whose targets lie outside it, such as SCH-79797/PAR1, are not recoverable (Supplementary S5). AlphaFold pockets are ligand-free and can be collapsed, and docking scores are approximate.

### Next steps

Direct experimental tests: GPR35 β-arrestin and Ca^2+^ functional assays and a focused GPCR counter-screen, using re-isolated or synthesized ligiamycin A.

## Methods

### Model

The architecture adapts the HitScreen sequence-based DTI framework [1]: an MPNN ligand encoder (PyTorch Geometric NNConv [9]); an ESM-2 protein encoder (facebook/esm2_t12_35M_UR50D, frozen [8]); bidirectional multi-head cross-attention fusion; and an MLP head. It was trained with binary cross-entropy plus an InfoNCE contrastive loss and same-ligand hard negatives. All code was written in Python, developed with the assistance of Claude Code, Anthropic’s agentic command-line coding tool.

### Data

BindingDB Kd subset [3]; active ≤ 100 nM, inactive ≥ 10,000 nM; 1,249 unique human target sequences.

### Screening and bias correction

Per-target sigmoid probabilities; background z-score across ≥ 6 molecules to remove frequent-hitter bias.

### Target identification

NCBI blastp against nr.

### Docking

Ligand 3D conformers from RDKit (ETKDGv3 + MMFF94 [10]); receptors from AlphaFold DB [7] (O14842, Q9HC97, P25021); CB-Dock2 [4] cavity-detection blind docking with AutoDock Vina scoring [5]; matched-pocket comparison versus positive (TAK-875, zaprinast, famotidine) and negative (glucose) controls.

## Supporting information

RESULTS

## Data and code availability

All code and data are available at https://github.com/docmartin95/hitscreen. See RESULTS_ligiamycin_GPR35.md and SUPPLEMENTARY.md. A tagged release archived to Zenodo will provide a citable DOI on publication.

