## Supplementary material for "A control-validated pan-proteome deep-learning pipeline nominates GPR35 as a candidate target of the orphan bacterial metabolite ligiamycin A": RESULTS

### Computational target prediction for the orphan bacterial metabolite ligiamycin A

#### Summary

Using a pan-proteome deep-learning drug–target interaction (DTI) model followed by bias-corrected ranking and structure-based docking with proper controls, we nominate **GPR35 (a class-A, lipid-sensing G-protein-coupled receptor)** as a candidate target of **ligiamycin A**, a 2022-described *Streptomyces/Achromobacter* co-culture metabolite (decalin-amino-maleimide) for which . The prediction is supported independently by (i) the model's bias-corrected
ranking and (ii) physics-based docking in which ligiamycin scores comparably to the known GPR35 agonist zaprinast at the agonist pocket. The result is a computational hypothesis requiring experimental confirmation.

#### Methods

- **Model.** Pan-proteome DTI model combining an MPNN ligand encoder (PyTorch Geometric), an ESM-2 protein language-model encoder (facebook/esm2_t12_35M_UR50D), bidirectional cross-attention fusion, and an MLP head, trained with BCE + InfoNCE contrastive loss and hard-negative sampling.
- **Training data.** BindingDB Kd subset (37,221 ligand–target pairs; 1,249 unique human target sequences). Trained on CPU; validation ROC-AUC ≈ 0.86.
- **Screening.** Ligiamycin A (SMILES O=C1NC(=O)C(N)=C1C(=O)C2(C)C3CCC(C)CC3C=CC2C, $C_{18}H_{24}N_2O_3$, structure derived from the published figure and RDKit-verified) was scored against all 1,249 unique targets.
- **Bias correction.** Raw model scores showed a "frequent-hitter" bias (a fixed set of heavily-measured proteins scored high for most ligands; confirmed by a losartan positive control that failed to rank AT1R first). Scores were re-ranked by per-target z-score relative to a background of 6 other molecules.
- **Docking.** CB-Dock2 (cavity-detection blind docking, AutoDock Vina) against AlphaFold models (FFAR1 O14842, GPR35 Q9HC97, histamine H2 P25021), each with a known-ligand positive control and a glucose negative control. Comparisons were made at the *matched* pocket (identified by the positive control's pose).

#### Results

### ML ranking

After background correction, ligiamycin A's standout targets were uniformly class-A GPCRs (BLAST-identified): **GPR35 (top hit, z = 1.55)**, bradykinin B1 receptor, free-fatty-acid receptor 1 (FFAR1/GPR40), somatostatin receptor 5, and histamine H2 — several of them lipid/peptide-sensing receptors, consistent with ligiamycin's lipophilic decalin scaffold. The specific receptor identity was unstable between raw and corrected rankings (the losartan control showed specific calls are unreliable), so the robust claim is **class-level (GPCR), not a single receptor**.

### Docking (matched-pocket, with controls)

| Receptor | Positive control @ true pocket | Ligiamycin @ same pocket | Glucose | Verdict |
|---|---|---|---|---|
| FFAR1 (O14842) | TAK-875 −7.1 (works) | −5.5 | −5.7 | Negative — not supported |
| **GPR35 (Q9HC97)** | **zaprinast −8.3 (works)** | **−8.1** | −5.5 | **Positive — corroborated** |
| Hist. H2 (P25021) | famotidine −5.8 (failed; collapsed pocket) | −7.6 (uninterpretable) | −5.5 | Inconclusive |

At GPR35's agonist pocket, ligiamycin (−8.1 kcal/mol) scored within 0.2 kcal/mol of the known agonist zaprinast (−8.3) and ~2.6 kcal/mol below the glucose non-binder floor (−5.5). Zaprinast (MW 271) and ligiamycin (MW 316) are similar in size, so the match is not attributable to the known Vina size/lipophilicity bias. Of three GPCRs tested, docking supported only GPR35 — the same receptor the ML model ranked first.

### Limitations

- Entirely computational; no experimental binding data.
- Model trained to ~0.86 validation ROC-AUC on a CPU-limited run; the losartan control showed the model resolves target *class* better than specific identity.
- AlphaFold receptor models are ligand-free; deep pockets (e.g. H2) can be collapsed, invalidating docking there.
- Docking scores are approximate; corroboration ≠ proof.

### Recommended experiments

1. **GPR35 functional assay** (β-arrestin recruitment or $Ca^{2+}$ flux; commercially available) to test agonism/antagonism by ligiamycin A.
2. **Broad GPCR panel screen** (e.g. β-arrestin gpcrMAX) to test the class-level hypothesis unbiasedly and catch the specific receptor.
3. Requires re-isolated or synthesized ligiamycin A (not commercially available).

### Reproducibility

#### Code files (pan_dti/ package)

| File | Role |
|---|---|
| data_pipeline.py | SMILES[illegible] molecular graph featurization (RDKit), DTIDataset, cold-start/random split, hard-negative sampling |
| dti_model.py | PanProteomeDTI: MPNN encoder + ESM-2 encoder + bidirectional cross-attention + MLP head |
| convert_bindingdb.py | Builds bindingdb_dataset.tab from the BindingDB Kd pull (active/inactive thresholds 100 nM / 10000 nM) |
| train.py | Training loop (BCE + InfoNCE), checkpointing to pan_dti_model.pt, CLI flags (--epochs, --max-rows, --batch-size, --smoke-test) |
| screen_sch79797.py | Score any molecule (--smiles, --name) vs all panel targets; optional NCBI BLAST (--blast); saves screen_results_.csv |
| background_rank.py | Frequent-hitter bias correction: per-target z-score vs background molecules |
| blast_top_zscore.py | BLAST-identify top background-corrected hits (--min-raw filters z-score artifacts) |
| check_panel_coverage.py | Checks whether a UniProt target is present in the training panel |

#### End-to-end workflow (commands actually run)

```
# 1. Build dataset (BindingDB Kd -> 37,221 pairs, 1,249 unique targets)
python -m pan_dti.convert_bindingdb

# 2. Smoke test the pipeline, then full training (ESM-2 weights auto-download)
python -m pan_dti.train --smoke-test
python -m pan_dti.train              # -> pan_dti_model.pt (val ROC-AUC ~0.86)

# 3. Screen the orphan compound
python -m pan_dti.screen_sch79797 \
```

```
  --smiles "O=C1NC(=O)C(N)=C1C(=O)C2(C)C3CCC(C)CC3C=CC2C" \
  --name "Ligiamycin_A" --blast --blast-top-n 8

# 4. Build a background set (re-run several molecules without --blast), then
#    bias-correct and identify the standout hits
python -m pan_dti.background_rank --molecule Ligiamycin_A
python -m pan_dti.blast_top_zscore --molecule Ligiamycin_A --top 6 --min-raw 0.3

# (controls used to validate the model: losartan, staurosporine, indirubin, etc.)
```

### Environment

Python with torch, torch_geometric, transformers, rdkit, pandas, scikit-learn, biopython, tqdm. ESM-2 checkpoint facebook/esm2_t12_35M_UR50D. Trained on CPU.

### Docking

Ligand 3D structures (ligiamycin A and all controls: TAK-875, glucose, zaprinast, famotidine) generated with RDKit (ETKDGv3 + MMFF94). Receptors: AlphaFold models O14842 (FFAR1), Q9HC97 (GPR35), P25021 (H2). Docking via CB-Dock2 (https://cadd.labshare.cn/cb-dock2/), AutoDock Vina scoring.

### Note on scope

The pan_dti/ package is a fresh pan-proteome model built for this study. An earlier, separate classical-ML pipeline also exists in the repo root (train_real_model.py, screen_sch79797.py, fetch_chembl_data.py, etc.) trained on a small kinase panel (toy targets + DAVIS); it is not used for the ligiamycin/GPR35 result above.
