## SUPPLEMENTARY DATA for "A control-validated pan-proteome deep-learning pipeline nominates GPR35 as a candidate target of the orphan bacterial metabolite ligiamycin A"

### Supplementary data — ligiamycin A target prediction

All scores are from the pan-proteome DTI model (predicted_prob_active, 0–1) or AutoDock Vina (kcal/mol) as indicated. Target identities are NCBI blastp assignments (e-value 0.0, ≥99% identity unless noted).

#### S1. Ligiamycin A — raw model ranking (top hits)

Top targets are uniformly class-A GPCRs (DRY / NPxxY motifs in sequences).

| Rank | Target (BLAST) | Class | Score |
|---|---|---|---|
| 1 | Neurotensin receptor 2 (NTSR2) | GPCR | 0.598 |
| 2–6 | Angiotensin II type-1 receptor (AGTR1) | GPCR | 0.59–0.60 |
| 7 | FLT3 | RTK | 0.578 |
| 8 | Histamine H4 receptor | GPCR | 0.566 |

Total predicted active (≥0.5): 128 / 1249.

#### S2. Ligiamycin A — background-corrected ranking (z-score) with BLAST IDs

Background = 6 other molecules; z = (score − bg_mean)/bg_std. Filtered to raw ≥ 0.3 to remove low-variance artifacts.

| z-score | raw | Target (BLAST) | Ligand class normally bound |
|---|---|---|---|
| 2.05 | 0.361 | (no BLAST hit — partial seq) | — |
| 1.55 | 0.440 | **GPR35** (orphan) | lipids / kynurenic acid |
| 1.48 | 0.463 | Bradykinin B1 receptor | peptide |
| 1.38 | 0.525 | Free fatty acid receptor 1 (FFAR1/GPR40) | fatty acids |
| 1.38 | 0.337 | Somatostatin receptor 5 (SSTR5) | peptide |
| 1.37 | 0.342 | Histamine H2 receptor | amine |

All credible (raw ≥ 0.3) standout hits are class-A GPCRs.

### S3. Model validation — control molecules (screen)

| Molecule | Known target(s) | Model's top hit(s) | Outcome |
|---|---|---|---|
| Staurosporine | pan-kinase | RSK1, Aurora B, RSK4 (kinases), ~0.75 | recovers kinase class □ |
| Honokiol | debated (EGFR/ FOXM1/Hsp27) | FLT3, KIT (RTKs), 0.84–0.86 | RTK class (consistent w/ EGFR hyp.) |
| Indirubin | CDK1/CDK5/ GSK-3β | FLT3, PARP1, KIT; all <0.5 (max 0.43) | did NOT recover CDKs (likely absent from Kd panel) |
| Losartan | AT1R (AGTR1) | PARP/kinase/KIT/NR ~0.85; AT1R only ~0.77 (rank ~12) | **fails to rank AT1R first □ frequent-hitter bias** |

The losartan result motivated the background-correction step (S2) and the conclusion that the model resolves target *class* more reliably than specific identity.

### S4. Docking — matched-pocket comparison (AutoDock Vina via CB-Dock2)

"True pocket" = pocket where the known-ligand positive control's best pose lands.

#### S4a. FFAR1 — crystal 4PHU (contains T4-lysozyme fusion)

| Ligand | Best Vina | Site of best pose |
|---|---|---|
| TAK-875 (positive) | −9.0 | real FFAR1 pocket (template-matched) |
| Ligiamycin A | −7.8 | T4-lysozyme fusion (artifact) |
| Glucose (negative) | −4.6 | T4-lysozyme fusion |

#### S4b. FFAR1 — AlphaFold O14842 (fusion-free)

| Ligand | @ true pocket (5,−4,−6) | Verdict |
|---|---|---|
| TAK-875 (positive) | −7.1 (best −8.8) | works |
| Ligiamycin A | −5.5 | **= glucose □ not supported** |
| Glucose (negative) | −5.7 | floor |

#### S4c. GPR35 — AlphaFold Q9HC97

| Ligand | @ true pocket C1 (10,11,−9) | Verdict |
|---|---|---|
| Zaprinast (positive) | −8.3 | works |
| **Ligiamycin A** | **−8.1** | **≈ agonist □ SUPPORTED** |
| Glucose (negative) | −5.5 | floor |

Zaprinast (MW 271) vs ligiamycin (MW 316) are similar size □ match not a size artifact.

#### S4d. Histamine H2 — AlphaFold P25021

| Ligand | Best Vina | Verdict |
|---|---|---|
| Famotidine (positive) | −5.8 (≈ glucose) | **control failed (collapsed pocket) □ inconclusive** |
| Ligiamycin A | −7.6 | uninterpretable |
| Glucose (negative) | −5.5 | floor |

### S5. Negative scope checks (compounds whose targets lie outside the panel)

| Compound | Reported target | Class | Model result |
|---|---|---|---|
| SCH-79797 | PAR1 | GPCR (not in Kd panel) | could not recover (panel gap) |
| Schweinfurthin A | OSBP/ORP4 | lipid-transfer protein | could not recover (panel gap) |

These illustrate the central limitation: the model can only rank targets present in the training panel (BindingDB Kd, 1,249 unique human sequences).
